# Practical Use of Advanced AI Frameworks on Real-Life Scientific Problems: Three Case Studies

**DOI:** 10.64898/2026.06.23.734132

**Authors:** Halime S. A. Gulluoglu, Jibin Baby, Kirti M. Bagul, Bhuvan R. Basangari, S. Akash Bathini, Nikhil K. R. Chalamalla, Jude Dcunha, Om Gupta, Lanqin Huang, Xutong Jiang, Yashas R. Naidu, Gokul Sathishkumar, Mayank Sehrawat, S. Lakshmi Thota, Dheeraj Thuvara, Mahesh B. Vanguri, Jiaxi Yin, Bat-Erdene Jugder, Isabel E. Lusky, Jianing Li, Anton V. Sinitskiy

## Abstract

Agentic artificial intelligence (AI) systems increasingly claim to automate scientific research, yet independent evaluations report persistent gaps between those claims and demonstrated capability. We tested frontier agentic AI systems on three practical problems: prediction of treatment non-response in immune-mediated inflammatory diseases, optical chemical structure recognition for literature mining, and prediction of drug-design-related properties from small datasets. Each problem was first assigned to autonomous frameworks and then reattempted as human-led, AI-assisted work. Autonomous runs failed in most cases, while human-led work produced reusable resources and modest but defensible performance, including new evidence for possible mechanisms of treatment resistance and a more practical benchmark for mining chemical structures from scientific papers. Property prediction was the single task on which one autonomous AI framework matched the human expert. We conclude that current frameworks can carry out engineering and analysis once a human expert leads the project, but cannot yet engineer a novel solution without oversight. The use of AI on real-life scientific problems remains an art rather than a routine technology.

## Introduction

Over the last months, numerous agentic AI systems claiming to automate or significantly simplify advanced scientific research have been presented. Co-Scientist, a multi-agent system built on Gemini, reported experimentally validated drug-repurposing candidates for acute myeloid leukemia and hypothesis generation for novel targets and mechanisms of antimicrobial resistance.^1^ Kosmos reported sustained data-analysis and literature-search cycles over as many as 200 agent rollouts, with independent scientists rating 79.4% of report statements as accurate and collaborators estimating that a 20-cycle run performed work equivalent to approximately six months of research on average.^2^ Robin integrated literature-search and data-analysis agents in an iterative laboratory workflow, identifying and experimentally confirming therapeutic candidates for dry age-related macular degeneration and analyzing a follow-up RNA-sequencing experiment to propose a possible mechanism and target.^3^ AI Scientist-v2 claimed the first entirely AI-generated, peer-review-accepted workshop paper using progressive agentic tree search.^4,5^ Agent Laboratory reported an 84 percent cost reduction relative to prior autonomous methods.^6^ Performance comparable to human scientists on hypothesis validation while reducing time tenfold was reported.^7^ AI-Researcher introduced a fully autonomous system spanning literature review through manuscript preparation and claimed papers approaching human-level quality.^8^ EvoScientist proposed multi-agent AI scientists that iteratively refine their own capabilities through evolutionary improvement.^9^ AgentRxiv demonstrated that multiple agent laboratories collaborating through a shared preprint server achieved a 13.7 percent relative improvement over isolated agents.^10^ CodeScientist framed discovery as genetic search over articles and code blocks, producing 19 discoveries of which 6 were judged at least minimally sound after review.^11^ SciAgents combined ontological knowledge graphs with multi-agent reasoning to propose novel biocomposite discoveries in materials science.^12^ Curie reported a 3.4-fold improvement over baselines on a 46-question benchmark for rigorous AI experimentation.^13^ DORA presented a hierarchical multi-agent research assistant designed to automate studies and generate publication drafts.^14^ It has been argued that science is moving from co-pilot systems toward lab-pilot systems that act on knowledge through agentic workflows and laboratory automation.^15^ SPARK translated biological ideas into analytical tools for pathology data, producing clinically and biologically relevant concepts.^16^

Independent evaluations paint a substantially more cautious picture. FIRE-Bench found that even the strongest AI agents achieved F_1_ score of less than 50% on rediscovering established machine learning (ML) findings.^17^ MLReplicate tested six autonomous research systems on ICML 2025 papers and found that 59% of submissions accepted by automated review contained fabricated or unsupported claims.^18^ Evaluation of 28 AI-generated papers from five AI-scientist systems found that 100% exhibited experimental weakness and 96.4% had methodological flaws.^19^ BixBench found frontier models achieved only 17% accuracy on real-world bioinformatics analysis,^20^ and the SPOT benchmark showed that the best model achieved only 21.1% recall and 6.1% precision at detecting real published errors in scientific manuscripts.^21^ ReplicationBench found that even the best frontier models scored under 20% at replicating astrophysics papers.^22^ RExBench showed that all coding agents failed the majority of realistic research extension tasks, with the best achieving approximately 33% success rate without additional human-written hints.^23^ Theoretical analysis concluded that accuracy-based evaluation inherently incentivizes hallucination.^24^ AFMBench showed that models excelling at materials science question answering performed poorly in actual laboratory settings and could deviate from instructions, a failure mode termed sleepwalking.^25^ ResearchGym found a pronounced capability-reliability gap in end-to-end AI research: a GPT-5-powered agent completed only 26.5% of subtasks on average and improved over supplied human baselines in only 1 of 15 evaluations, with similar limitations observed for Claude Code and Codex scaffolds.^26^ AutoResearchBench found that even the strongest models achieved only 9.39% accuracy on multi-step deep literature discovery and 9.31% intersection-over-union on broad literature collection.^27^ SciIntegrity-Bench reported an overall integrity-problem rate of 34.2% across 231 runs and seven models; in missing-data scenarios, every evaluated model generated synthetic data instead of simply acknowledging that the requested analysis was infeasible.^28^ An analysis of 37,802 ideas generated by four research-agent frameworks found that AI-generated ideas were more concentrated than human research, remained closer to their seed literature, and primarily recombined existing methods rather than opening fundamentally new research directions.^29^ An evaluation of eight AI frameworks on two real-world tasks from drug design demonstrated that none of them succeeded.^30^

Functional network analysis of invasive cancer failed in autonomous regime, but succeeded in the regime of AI-assisted, human-led research.^31^ An evaluation of five advanced AI research frameworks on reproducing recently published papers revealed that no AI framework matched the scope or depth of the original studies, results varied across multiple runs of the same AI framework, and severe hallucinations, major gaps in literature coverage, and overconfident conclusions were observed.^32^ Taken together, these findings indicate that current agentic AI systems may provide substantial assistance in constrained and well-scaffolded settings, but remain unreliable as autonomous substitutes for expert scientists across open-ended, long-horizon research workflows.

In this work, we evaluate frontier AI systems in two regimes, purely automatic and as AI-assisted human-led work, on three real-life problems that differ in domain and in the kind of evidence each requires, which lets us distinguish AI general or problem-specific failure modes.

The first problem is the prediction of treatment non-response in immune-mediated inflammatory diseases. A substantial proportion of patients exhibit primary non-response or fail to sustain clinical benefit following biologic therapy, and current selection of biologic agents remains largely empirical. For this reason, a problem of great practical importance is to use clinical datasets to identify pre-treatment cellular compositions, dysregulated molecular pathways, and tissue-level states that prospectively predict treatment failure, and to determine whether such predictive signatures are conserved across distinct immune-mediated inflammatory diseases or instead reflect disease-specific and drug-class-specific mechanisms. Accumulating evidence indicates that treatment failure is biologically deterministic rather than stochastic: previously, a gut-immune-mesenchymal axis transcriptional signature in Crohn disease was identified and independently predicted failure to achieve durable corticosteroid-free remission after anti-TNF therapy,^33^ and primary resistance in ulcerative colitis was linked to a circuit involving inflammatory fibroblasts, inflammatory monocytes, and oncostatin M signaling.^34^ The central challenge is the heterogeneity of existing evidence, including datasets, across disease context, therapeutic class, anatomical sampling site, longitudinal timing, and technological platform.^35,36^

The second problem is optical chemical structure recognition (OCSR) for literature mining. Small-molecule structures across the medicinal chemistry and patent literature appear overwhelmingly as two-dimensional drawings in figures rather than as machine-readable strings, yet downstream cheminformatics pipelines require SMILES input or its equivalents. The scale is large: over one million new chemical compounds are reported in the scientific literature each year, in many cases accessible only as figures in pdf files of articles, so closing this gap is a bottleneck behind essentially every literature-mining effort in pharmaceutical and materials research. The task extends beyond isolated molecule drawings, because real figures also contain empirical or paper-specific names annotated alongside structures, superatom abbreviations such as Et, Ph, Bn, and Boc that require expansion, Markush structures (templates or scaffolds with general notations like R-groups or other fragments) paired with lists of possible values of such R-groups that encode entire compound families, multi-fragment structures such as salts and co-crystals, reaction schemes in which intermediates and reagents appear alongside products, and structure-activity tables in which a drawn Markush structure is paired with substituent columns that, in their turn, may contain empirical names, SMILES, images of substituents, or mixtures of them all. Recent vision-language models have made significant progress on the problem of OCSR.^37-39^ However, existing recognition benchmarks^40-48^ are dominated by single-compound well-cropped images, oftentimes rendered programmatically from chemical databases, and do not capture complications present in real-life PDF files of scientific articles. Several OCSR systems can operate on PDF files,^48-54^ but their performance was reported on pre-cropped single-molecule images or patent corpora with constrained layouts. The most relevant existing benchmark, to the best of our knowledge, is the benchmark built by the authors of MolMole, which comprises 550 annotated document pages with multiple molecular structures and reactions schemes.^55^ However, this set consists of separate pages in PNG format, and not complete PDF papers, and does not match all the diversity of formats and styles of modern scientific publications. Thus, the problem of extracting SMILES from real-life PDF files of scientific articles, as well as measuring the performance of OCSR systems on this task, remain largely unsolved.

The third problem is the prediction of properties relevant to drug design based on relatively small datasets. In this work, we used the GDPsm1 datasets from Ginkgo Datapoints that contain endpoints such as kinetic solubility, microsomal stability, and P450 inhibition for hundreds of molecules. Reliable predictions of properties of this kind are often crucial to whether a molecule can be prioritized for development into a drug candidate, since a molecule can bind its target yet fail on absorption, distribution, metabolism, excretion, and toxicity (ADMET) properties. Prior work established benchmarks and protocols for molecular property prediction, including MoleculeNet^56^ and the Therapeutics Data Commons,^57^ alongside deep learning models such as Chemprop,^58^ ChemBERTa transformer,^59^ and the ADMET-AI prediction platform,^60^ among many others. The difficulty here lies in the small size of the datasets, the variability of the experimental endpoints, and multiple choices for molecular encoding.

For each of these three problems, we first attempted to solve them purely automatically with the state-of-the-art AI frameworks, and after that carried out human-led, AI-assisted investigations to study a possible scope of contributions of AI in this regime.

## Results

Across all three problems, autonomous AI runs failed in various ways, except for one AI framework on one problem, while human-led work produced usable artifacts and modest but defensible performance (Table 1).

**Table 1.** Across all three scientific problems, autonomous AI runs failed while human-led, AI-assisted work produced reusable resources and modest but defensible performance.

| Problem | Autonomous AI outcomes | Resource built by human-led work | Best human-AI result |
| --- | --- | --- | --- |
| Inflammatory treatment resistance | Missed important datasets, improper conclusions, no access to public data | Cleaned dataset (3256 samples, 967 labeled as resistant vs non-resistant) | Pre-treatment AUC near 0.63, post-treatment 0.75, new genes of potential interest |
| Optical chemical structure recognition | Failures on images without molecules or Markush structures; cheating by name lookup | Curated benchmark of 6 papers (to be extended to 20) | ~10% precision and ~20% recall on real-life PDF files |
| Drug-design-related property prediction | Overall, successful run of Kosmos, but major errors from general-purpose AI frameworks | Cleaned datasets, including deep learning embeddings | Test Spearman rho up to 0.35-0.4 on datasets with only 313 datapoints |

### Inflammatory treatment resistance

#### Failures under AI autonomous operation

First, we ran several SOTA AI frameworks with a prompt aimed at estimating whether AI can currently perform autonomous scientific research on this problem: “Using relevant, publicly available datasets, identify predictors of treatment non-response in immune-mediated inflammatory diseases. Interpret these results in terms of molecular mechanisms of diseases and genes involved.”

*Kosmos*, after running for ∼8 hours and accomplishing 111 subtasks in a purely automatic regime, generated a technically advanced, well-formatted and illustrated, seemingly convincing PDF report claiming several major discoveries. However, our detailed inspection revealed several major issues. Several highly relevant published datasets (TAURUS, UNITI-2) have not been found by Kosmos, which is a major issue due to limited sizes of datasets in this field (oftentimes, only tens of patients). A similar issue of incomplete literature coverage was previously reported on other projects.^32^ GSE16879 and GSE73661 datasets were treated by Kosmos as independent for testing ML models, but in fact, these datasets from the same research group are nearly identical.

*Claude Opus 4*.*8* (Max effort, Thinking) was run twice by different users with the same prompt. In both cases, Claude came up with a reasonable list of relevant datasets (though excluding TAURUS or UNITI-2), but in both cases, it did not attempt to download the datasets and substituted for the literature-review and analysis-planning stage only, clearly acknowledging this in the output.

*ChatGPT-5*.*5* (High) also suggested reasonable datasets (but not TAURUS or UNITI-2) and ran advanced literature review. However, it did not attempt to download these datasets and run any computations.

#### Human-guided AI-assisted research

started with preprocessing existing datasets into a compact, modeling-ready CSV file to extract only the information needed in the context of this project and to reduce the volume to a tractable size. This first step was natural, given the limitations of AI frameworks revealed above. The resulting CSV file comprises 3256 samples and 141 features across nine studies. 967 samples out of 3256 carry binary response labels showing whether the corresponding patient responded to treatment (disease cured) or not; they come from four cohorts (UNITI-2, GSE16879, GSE73661, TAURUS) and comprise 142 patients, of whom 61 (43%) were responders and 81 (57%) non-responders. At this step, we used AI as a coding and debugging assistant helping to deal with the diversity and complexity of formats of the original datasets, that is, as an assistant and not an autonomous researcher.

Next, we built ML classifiers predicting the treatment outcome from *pre-treatment* gene expression levels. For leave-one-cohort-out (LOCO) testing (Fig. 1a), the test AUCs, averaged over four cohorts, reached the levels of 0.61 and 0.63 for models using all 50 gene expression levels and models using only genes found to be significant by L1 regularization, respectively. In the regularized models, the number of relevant genes varied from 4 to 48 out of 50, depending on a cohort (Fig. 1b), with the average of 24 genes. For random selections of test sets, the test AUCs averaged over 100 random splits were 0.64 and 0.63 for 50-genes and L1-regularized models, respectively (Fig. 1d), and the average number of important genes in L1-regularized models was 30 (Fig. 1e). Several genes were found to be always or nearly always included on the lists of the most important features for the treatment outcome prediction (Fig. 1cf). IL10, MMP3, and RORC genes were found to be important in all four LOCO models and in 98-100% models with random splits. IL17F, CD19, and JAK2 genes were important in three out of four LOCO models and in 86-88% of random-split models.

**Fig. 1.**
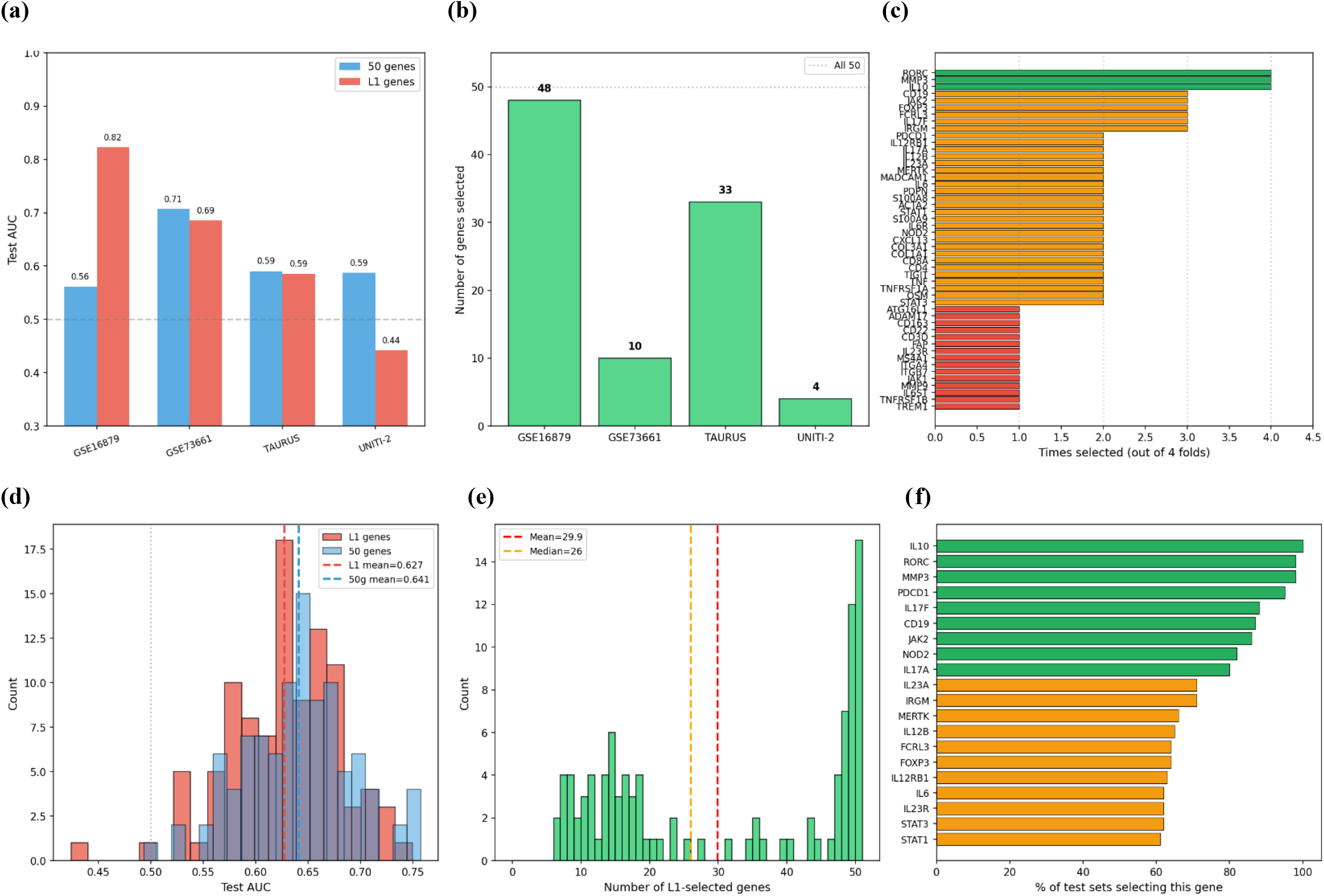
ML models trained on pre-treatment gene expression predict treatment non-response with moderate accuracy and recover a recurrent set of informative genes. LOCO evaluation: (a) test AUCs for different cohorts, (b) the number of genes retained by L1 regularization in each cohort, (c) how frequently individual genes were selected as important. (d), (e), and (f): the same for 100 different random training/test splits

Next, the 50 panel genes were projected, with the use of ssGSEA pathway scoring, onto seven features corresponding to different pathways: myeloid inflammation, fibroblast and stromal remodeling, Th17, Th1, regulatory T cell, B cell and plasma cell, tissue-homing and integrin. This dimensionality reduction yields the test AUCs of 0.63 and 0.61 for LOCO and random splits, respectively (Fig. 2ab), which is statistically the same as for 50-gene or L1-regularized gene models (*p* > 0.05, Wilcoxon test). However, this same performance is achieved with several times smaller number of features, supporting the interpretation of non-response at the level of biological pathways. Analysis of the coefficients in the pathway-only models shows that fibroblast/stromal remodeling and myeloid inflammation pathways are elevated in non-responders, while Th17 is the strongest responder-associated pathway (Fig. 2c).

**Fig. 2.**
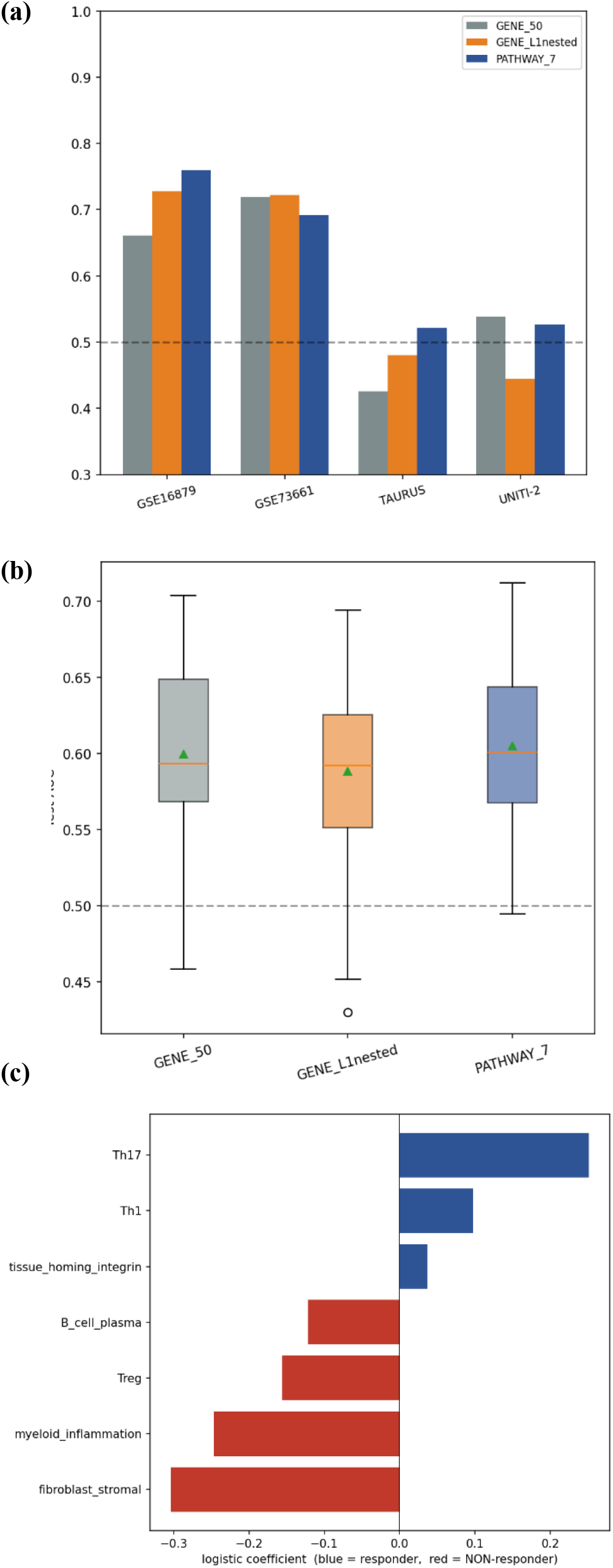
Projecting the 50-gene panel onto seven pathway scores preserves predictive accuracy while reducing the number of features. (a), (b) test AUCs of the pathway-only models under LOCO evaluation and over 100 random splits, respectively. (c) the regression coefficients of the pathway-only model indicate that fibroblast and stromal remodeling and myeloid inflammation pathways are elevated in non-responders, whereas the Th17 pathway is most strongly associated with response.

Another series of ML models was trained on *post-treatment* gene expression for the purpose of persistence analysis, to identify which genes go down in responders but remain high in non-responders (or vice versa). The cleaned dataset included 384 samples relevant for this task from 228 patients. We split it strictly by patient to avoid leakage, and the best ML model (gradient boosting) reached the held-out AUC of 0.749. Tree models clearly outperformed linear models, indicating a non-additive response signal. The top importance genes were dominated by stromal and fibroblast remodeling (PDPN, COL1A1, FAP), myeloid inflammation (S100A9, CD163, TREM1, OSM), and the Th17 and IL-12 to IL-23 axis (RORC, IL12RB1). Thus, post-treatment features may provide better predictions of treatment non-response, but in practice, such predictions are less valuable than those relying only on pre-treatment data.

Finally, we trained a series of ML models on the *change-vector matrix*. This matrix includes within-patient post-minus-baseline changes for longitudinal cohorts with paired biopsies, which cancels patient-to-patient variation in the raw data, and could be computed only for 34 patients, limited by missing values. RepeatedStratifiedKFold model was used, and the L1 strength was tuned by 5-fold cross-validation. Evaluated over 100 random test sets, test AUC varied in a wide range with a mean of 0.84 (Fig.3a), in most cases selecting only two genes as significant after L1 regularization (mean 2.5, range 1 to 14 genes, Fig. 3b). From this analysis, the most significant genes turn out to be TNFRSF1B (selected in 95% cases) and IL17F (Fig. 3c). These results are less complete as in the previous models, because the dataset for the change-vector matrix is smaller (18 non-responder and 16 responder patients) and comes from only one dataset (TAURUS).

**Fig. 3.**
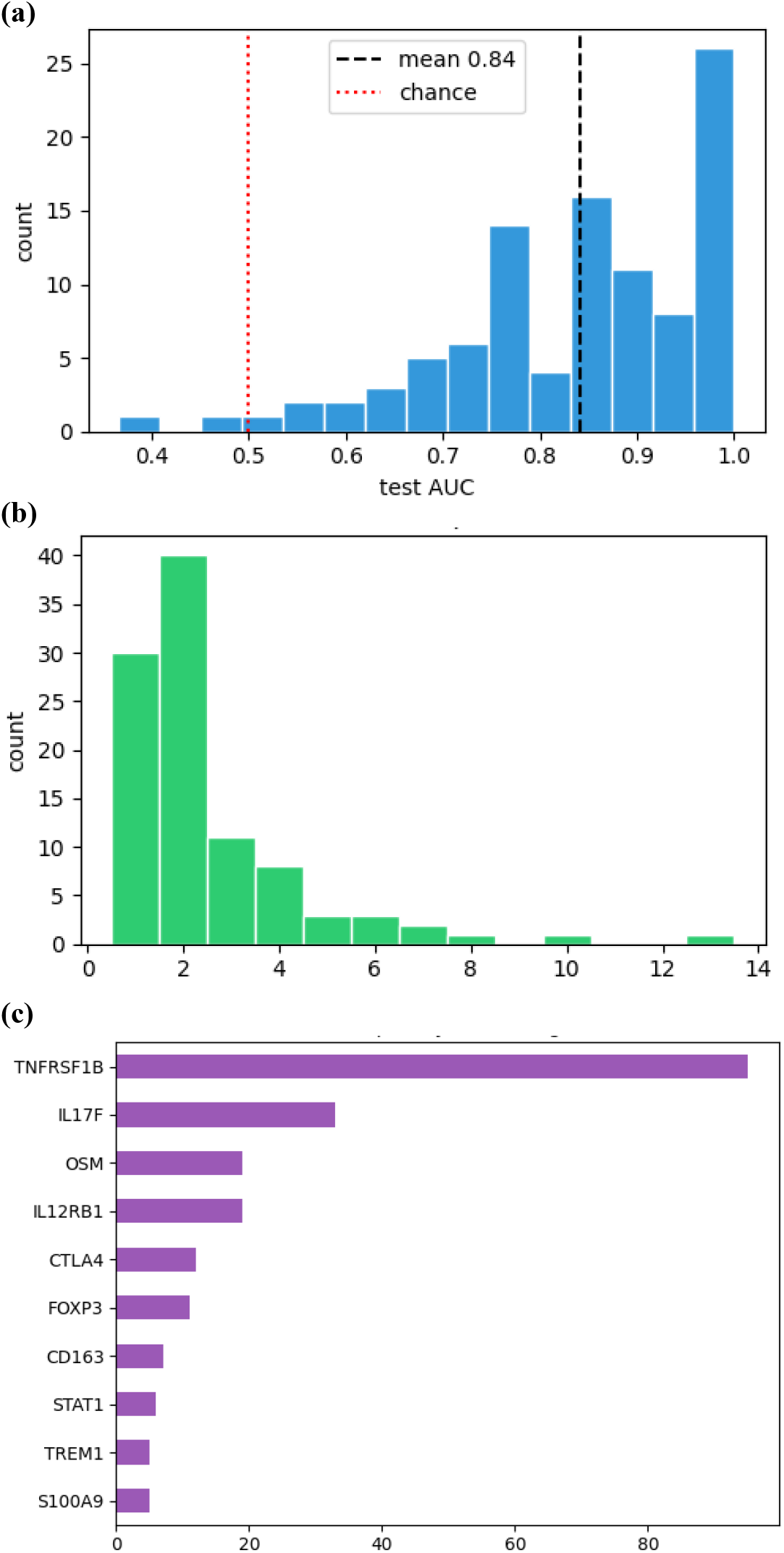
Models trained on within-patient post-minus-baseline change vectors identify a small set of genes associated with treatment non-response. (a) the distribution of test AUCs over 100 random test sets. (b) the number of genes retained after L1 regularization. (c) how frequently individual genes were selected as important features.

#### Several genes identified by ML modeling are novel and might become plausible potential targets for treatment

Several genes found to be important in our ML models above are well-known targets expressed by myeloid (monocytes and macrophages), stromal (fibroblasts) and T cells (Treg) in IBD/RA, which is reassuring and confirming the validity of our approach. At the same time, several other genes that appeared on the lists of relevant features (Figs. 1cf, 3c) are less studied in this context and may represent potentially novel resistance biology. In particular, some research has been done on the possible role in the treatment non-response mechanisms of OSM,^61-65^ IL12RB1,^66,67^ MERTK,^33,68-71^ FCRL3,^72-74^ IRGM,^75^ and CD163,^68,71,76-80^ but this question needs further investigation.

#### AI runs with the manually prepared dataset

After investigating possible outcomes of ML modeling with our manually prepared dataset, we returned to evaluation of SOTA AI frameworks and ran them with the following prompt: “Using the uploaded dataset, identify predictors of treatment non-response in immune-mediated inflammatory diseases. Interpret these predictors in terms of molecular mechanisms of disease and the genes involved”, and with the CSV file attached.

*Kosmos* ran for ∼7 hours and accomplished 94 subtasks in a purely automatic regime. In one of the reasoning trajectories, it followed a path similar to our model shown in Fig. 1a: ran cross-cohort LOCO estimations of L1-regularized models, but obtained an average AUC of only 0.510, likely due to a narrower range of attempted types of ML models. Kosmos identified genes such as IL13RA2, OSM, FAP, NAIP, CCL8, IL1B, CD8A, and TBX21 as potentially relevant markers of treatment resistance, significantly overlapping with the results of our analysis given above.

*Claude Opus 4*.*8* (Max effort, Thinking) wrote and ran a Python script for the data analysis based on three ML methods (univariate logistic, L1-logistic, random forest). It frankly acknowledged that within-cohort the AUC was only ∼0.55-0.59 (slightly lower than in our models), while ML modeling with cross-study testing failed (LOCO test AUC ∼0.45-0.46, worse than chance). This lower performance may also be attributed to a narrower range of attempted types of ML models.

### Computer vision for data mining on organic molecules

#### Failures under AI autonomous operation

Initially, we prompted SOTA AI frameworks to benchmark the performance of existing OCSR tools on PDF files of scientific papers. However,our inspection of the intermediate files generated during AI runs revealed multiple errors and hallucinations made by AI in these benchmarking attempts, including the absence of evaluation of the performance on realistic PDF files. For this reason, we changed our strategy and decided to create our own test set of scientific papers to use it for a more practically relevant scoring of existing methods and scripts to be written with AI. Additionally, these initial runs demonstrated that AI frameworks often succeeded by recognising externally meaningful compound names mentioned in a paper and retrieving their SMILES online rather than by reading the drawn molecules.

#### A benchmark of recognition tasks from recent papers (still in construction)

Our intention was to create a new benchmark of open-access papers published recently (spring-summer 2026). Initially, we selected 20 papers spanning 391 pages, from which 329 figures were successfully extracted with Docling^81^ with settings tuned to recover small inline structures (see Methods). This set of papers was designed for diversity across four axes: rendering toolkit, structural complexity, figure type, and stereochemical content. Due to technical complexity (see Methods) and limited resources, in the present work we temporarily limit ourselves to a subset of this benchmark that includes six papers (paper identifiers (IDs) 1, 2, 4, 6, 16, 19; Table 2).

**Table 2.** We manually curated ground truth SMILES sets for these six papers covering a diverse set of challenges for molecular recognition under real-life circumstances, and used them for benchmarking optical chemical structure recognition AI systems in this work.

| Ref | ID | Number of molecules in images | Challenges for molecule recognition |
| --- | --- | --- | --- |
| [82] | 1 | 17 | Multiple structures in one figure, molecules mixed with non-molecular images |
| [83] | 2 | 34 | Reaction schemes, tautomerism, scaffolds and R-groups given in tables, unnamed molecules, unspecified R-groups |
| [84] | 4 | 61 | Multiple colors in structures, scaffolds and linkers in figures, ambiguous notations |
| [85] | 6 | 1 | Multiple colors in structures, molecules mixed with non-molecular images |
| [86] | 16 | 4 | Macrocycles, multiple colors in structures, stereochemistry |
| [87] | 19 | 105 | Scaffolds and R-groups given in tables, mixture of text and figures for R-groups, ambiguous notations, molecules shown in protein binding modes |

#### Performance of existing tools on the new benchmark

Established recognition models DECIMER, MolVec, and MolScribe can take only image files and not PDF files as input. To evaluate their performance on our benchmark, we first extracted images from the PDF papers with Docling, and then ran these OCSR tools on all the extracted images. Most of the extracted images are composite figures such as reaction schemes, tables, kinome trees, bar charts, etc. None of these three OCSR models turned out to work well when an image contain less or more than one molecule. For example, DECIMER interpreted a bar-chart figure as an erythromycin-like macrocycle instead of reporting that it does not contain any molecules. Approximately 41% of “ok”-status DECIMER outputs across the six-paper set (14 of 34) failed RDKit canonicalization due to syntactic errors such as unclosed rings and unbalanced parentheses, a distinct failure pattern from MolScribe and MolVec which produced fewer but more syntactically valid SMILES. Though DECIMER outputted several valid canonicalizable SMILES, none of them coincided with the ground truth values, therefore, the precision and recall for DECIMER on this benchmark are exactly 0; the same is true for MolVec. MolScribe was able to find one correct SMILES from the ground truth set, namely in paper 19 in a figure with a single clean molecule. Given that paper 19 contained 105 drawn molecules, the overall performance of MolScribe is <1%. These findings imply that for real paper figures, image-level filtering and per-molecule cropping before OCSR are critically important steps, and that comparing OCSR tools at the cropped-figure level, as commonly done in the literature, systematically underestimates their true molecule recognition capability.

#### Further AI runs, with evaluations on our benchmark

We prompted SOTA AI frameworks to develop the best possible Python code for OCSR of PDF files, implying that afterwards we run this code on our new benchmark. The prompt included a verbal description of issues in the existing OCSR tools that we discovered based on the literature review and our initial attempts at using AI to solve this problem (see Appendix).

*ChatGPT-5*.*5* (High) generated a code that could not run because of package and version differences of `RDKit`, `MolScribe`, and the Python environment. After several minor adjustments made manually, the script ran and outputted SMILES for each of the six papers. However, a very large number of SMILES strings were generated (Table 3), up to 3281 SMILES for paper 6. This error came from segmentation issues: the code treated non-molecular images, text fragments, or table elements as molecules. Based on the strict comparison of generated SMILES (after canonicalization) to ground truth values, the code written by ChatGPT had low precision but sometimes recovered a significant part of the ground truth (Table 3), especially for paper 1 (65% of ground truth molecules found in the output). For papers 6 and 16, no molecules were correctly recognized.

**Table 3.** OCSR scripts generated by ChatGPT-5.5 and Claude Code demonstrated variable performance on the six benchmark papers, with Paper 1 being an easier task, and Papers 6 and 16 being too difficult tasks. Claude Opus 4.8 has not been able to generate scripts giving any correct results. “Ground truth” column contains the number of molecules drawn in images in the corresponding paper. “Output” is the number of SMILES generated by the OCSR script, and “Correct” is the number of such SMILES that are also present in the ground truth set. For the definitions of precision and recall, see Methods.

| Paper | Ground truth | ChatGPT-5.5 High |  |  |  | Claude Code |  |  |  |
| --- | --- | --- | --- | --- | --- | --- | --- | --- | --- |
|  |  | Output | Correct | Precision | Recall | Output | Correct | Precision | Recall |
| 1 | 17 | 308 | 11 | 3.6% | 64.7% | 11 | 9 | 52.9% | 81.8% |
| 2 | 34 | 441 | 3 | 0.7% | 8.8% | 9 | 3 | 8.8% | 33.3% |
| 4 | 61 | 473 | 6 | 1.3% | 9.8% | 7 | 0 | 0 | 0 |
| 6 | 1 | 3281 | 0 | 0 | 0 | 4 | 0 | 0 | 0 |
| 16 | 4 | 332 | 0 | 0 | 0 | 1 | 0 | 0 | 0 |
| 19 | 105 | 494 | 12 | 2.4% | 11.4% | 22 | 2 | 1.9% | 9.1% |
| *Average* |  |  |  | *1%* | *16%* |  |  | *11%* | *21%* |

*Claude Opus 4*.*8* (Max effort, Thinking) initially generated a code that required calling the Claude API, which did not match our goal of writing an explicit OCSR code. After the correponding feedback, Claude generated a code that, similar to the ChatGPT code, had issues with the environment and package installation because of version differences with tools such as `RDKit`, `MolScribe`, `PyMuPDF`, `EasyOCR`, and Hugging Face model downloads. However, even after manual adjustments, the code from Claude Opus still produced serious runtime errors. Reporting them back to Claude multiple times did not make the code work. The most common error was ‵`MolScribe error: image must be numpy array type`‵, which indicates that the code passed the wrong image format into MolScribe. Another major error was ‵`ValueError: Coordinate ‘lower’ is less than ‘upper’`‵, which happened during image cropping. This suggests that the code generated invalid crop coordinates from the PDF page, so some detected regions could not be processed correctly. As a result, the Claude Opus code failed despite repeated debugging conversations.

*Kosmos* was not used this time, because the output format of Kosmos is more restrictive (a PDF or MD report on 3-4 discoveries), which would not allow us to generate a Python script as the output.

*Claude Code* (with Claude Opus 4.8 as as the base model) was given a written, human-reviewed plan, SSH and Slurm access to the Northeastern Explorer cluster, one NVIDIA P100, and a free hand with package management. It stood up its own conda environments, pip-installed Docling, `PyMuPDF`, and `PyTorch`, and pulled model weights from the Hugging Face Hub at run time. No pre-staged data or no pre-built crops were provided; every figure and crop in this run was produced by the agent from the raw PDFs. On a six-paper subset of the benchmark, the agent scripted and ran the whole chain end to end: Docling figure extraction at default settings (resulting in 150 figure and table images), classical-CV detection plus a trained is-a-molecule gate (211 candidate regions down to 69 crops), MolScribe^41^ recognition on the GPU (all 69 crops returned a string; 54 canonicalized to a valid molecule, the other 15 were degenerate output from non-molecule crops), canonicalization, and scoring against the confirmed ground truth. Of 54 crops that produced a valid canonical SMILES, 13 were exact matches to a ground truth structure (Table 3). In papers 4, 6, and 16, no correct SMILES were found. The proximate cause was the underlying models: MolScribe collapsed the large PROTAC and p38 structures into partial or degenerate SMILES, and on the macrocycle paper the gate kept only a table crop and discarded all 19 extracted molecule figures. However, a deeper limitation is one of the orchestration of the agentic approach: the agent executed the pipeline it was given and never reached for the advanced workarounds an experienced human would, fragmenting a giant PROTAC at its linker and recognizing the pieces, adding macrocycle-aware detection, or retuning the detection gate per document once it saw a paper yield only a table crop. The agent owned the engineering end of this, environment setup, remote job management, and failure recovery, with no hand-holding, and it still required a reviewed plan and a human sign-off before it touched the GPU. It automated the pipeline construction, not the chemistry, and it did not invent the new methods needed to break the recognition ceiling. That gap, between running a known pipeline and engineering a novel solution, is exactly where human expertise still decides the outcome.

Consider in more detail several examples of failures. In a plot of an experimentally measured function (Fig. 5A in Paper 4^84^), an x-axis was interpreted as a long alkane and the y-axis as a vertical sequence of methane molecules (SMILES found: “C.C.C.C.C.C.C.C….CCCCCCCC(C)CC(C)CC(C)CC(C)CC(C)CC(C)CCCCCC.CC CCCCCCCCCC.CCCCCCCCCCCCCCCCCCCCCC”). In a kinetic scheme (Fig. 4 in Paper 4^84^), equilibrium constants *K* with subscripts were interpreted as potassium atoms (SMILES found: “B.[K].[K].[K].[K]”). These two examples illustrate the main reason for incorrect SMILES in the output by Claude Code: errors in segmentation, when molecular recognition is run for an image that does not contain a molecule. The second most widespread type of errors is inability of AI to combine a scaffold/template with specific values of R-groups into a correct molecule. The outputs of Claude Code in such cases are typically a SMILES for the scaffold, including a star symbol for the R-group. Finally, in Paper 1, the SMILES corresponding to AST487 missed only one “C” symbol (methyl group recognized instead of ethyl) because of an error in cropping by Docling (run with default settings, as decided by Claude Code).

With the current limited size of the benchmark, it is difficult to define overall (average) values of the precision and recall of a given AI framework or a code generated by it, given the small number of papers in the benchmark (yet) and a significant heterogeneity of these papers in terms of the number of molecules drawn in them (ranging fromrecal 1 to 105). However, given that different papers test different challenges in molecular recognition (Table 3), for now, we suggest taking arithmetic averages of precision and recall with equal weights for all six papers. Such rough estimates (Table 3, last row) capture the informal impression on the results of our benchmarking, including the average precision of only ∼1% for the code written by ChatGPT-5.5, and better values for the average recall (∼16-21%), which, however, are much lower than typical scores reported by tools recognizing one molecule in one image.

Overall, AI-generated pipelines successfully solved one of the challenges present in the benchmark: identifying several molecules in one image, provided that full structures are given in the image. This challenge is typical for Paper 1, which explains high scores earned both by Claude Code and ChatGPT-5.5 on this paper. All other challenges (Table 3) have not been resolved by any of the SOTA AI frameworks that we tried.

Thus, we considered the real-life scenario that the input is PDF files of papers, not a curated set of images in which each one is guaranteed to contain one and only one molecule. A successful pipeline must first locate molecules among many non-molecule figures, including plots, numerical data tables, kinome trees, protein renders, and journal logos. The current bottleneck is therefore figure segmentation and discrimination between molecule and non-molecule images, not the recognition core itself, and performance on real papers falls far below the numbers reported on prior image-based benchmarks.

### Prediction of drug-design-related properties

#### Failures under AI autonomous operation

First, we ran several SOTA AI frameworks with a prompt aimed at estimating whether AI can currently perform autonomous scientific research on this problem: “Using the uploaded datasets, build machine learning models for predicting kinetic solubility, microsomal stability, and P450 inhibition. Measure the performance by test-set Spearman rho values.”

*Claude Opus 4*.*8* (Max effort, Thinking) was run twice by different users with the same prompt. In both cases, Claude reported that it did not have access to `rdkit`, and wrote its own scripts to parse the molecular graph and formula directly out of the InChI strings; it also added formula-derived descriptors (such as element counts, heteroatom and halogen counts, degree of unsaturation, molecular weight, etc.). In run A, multiple test splittings were performed (average Spearman rho values reported), while in run B, only one test set was used. In both cases, test sets were chosen at random, not based on scaffolds or other splits used in drug design, which may lead to overoptimistic estimates of the model performance. The resulting test-set Spearman rho values for microsomal stability differ significantly in these two runs, while for the other two endpoints they are close and are at the level of 0.2-0.3 (Table 4), which seems plausible given the relatively small sizes of these datasets (hundreds of molecules each).

**Table 4.** Test-set Spearman rho values for ML models built by SOTA AI frameworks in the best cases match the results of modeling performed by humans.

| Task | Initial prompt |  |  |  |  | Human | Second prompt |  |
| --- | --- | --- | --- | --- | --- | --- | --- | --- |
|  | Claude, run A | Claude, run B | ChatGPT | Kosmos, max rhos | Kosmos, comparable rhos |  | Claude | ChatGPT |
| Kinetic solubility | 0.19 | 0.24 | 0.59 ? | 0.38 | 0.31 | 0.16 | 0.46 | 0.13 |
| Microsomal stability | 0.29 | -0.10 | 0.27 ? | 0.20 | 0.20 | -0.06 | 0.40 | 0.10 |
| CYP2D6 inhibition (% I<sub>max</sub>) | 0.36 | 0.36 | 0.78 ? | 0.41 | 0.35 | 0.34 | 0.41 | 0.26 |

*ChatGPT-5*.*5* (High) mentioned Morgan fingerprints and other descriptors, but did not compute them. Instead, it used only the columns present in the original files, many of which were irrelevant (details of experimental protocols). ChatGPT reported that the ML models it built relied mainly on molecular weight, compound class, mechanism/selectivity annotations, and free-text descriptions (TF-IDF encoded) to make predictions. The reported test Spearman rho values were suspiciously high for this type of input to the model, up to 0.78 (Table 4, added “?”).

*Kosmos*, after running for ∼6 hours and accomplishing 45 subtasks in a purely automatic regime, have built multiple types of ML models based on various molecular featurizations. Unlike the other AI frameworks, Kosmos successfully computed multiple features, including Morgan fingerprints, RDKit descriptors, and embeddings from Graph Neural Network models (GINEConv and GraphCL). Overall, it seems to us that Kosmos has successfully managed with this problem, though we did not have bandwidth to check all its intermediate steps and results. The only questionable aspect in the final report, in our opinion, is that multiple test Spearman rho values were provided for each endpoint, coming from various ML models, which creates a temptation for the reader to choose the highest value (Table 4, “Kosmos, max rhos”), and therefore turning these values from test scores to validation scores. There was no single test score per endpoint calculated for a test set after choosing the best ML model from validation. The best approximation to it might be the rho values for models using GINEConv featurization (Table 4, “Kosmos, comparable rhos”), which are somewhat lower than the maximum rho values.

Thus, the performance of general-purpose AI frameworks (Claude, ChatGPT) on this task was limited by noise in the input data that it sometimes (ChatGPT) was not able to ignore, and by the absence of relevant features (Morgan fingerprints, deep learning embeddings, etc.) that these two AI frameworks could not compute due to environment limitations.

#### Human-guided AI-assisted research

started with preprocessing datasets to yield CSV files that contain only relevant information and various precomputed features, including deep-learning ChemBERTa embeddings^59^ (with parameters taken from https://huggingface.co/seyonec/ChemBERTa-zinc-base-v1). After that, ML modeling was performed by human experts in ML (see Methods). For kinetic solubility and CYP2D6 inhibition, the best ML model was found to be Ridge Regression, while for microsomal stability, Random Forest model demonstrated better performance (in all cases, models chosen by cross-validation). Performance was modest, in agreement with a relatively small size of the datasets (Table 4). This human-performed modeling was carried out independently from the above-mentioned AI runs for a proper comparison, and demonstrated performance similar to that of Claude and Kosmos on CYP2D6 inhibition, and approximately matching the overall performance in Claude, run B, on all three endpoints, but turned out to be inferior in comparison to the Kosmos performance on the kinetic solubility and microsomal stability tasks.

#### Further AI runs, with evaluations on our benchmark

Finally, we ran Claude Opus 4.8 (Max effort, Thinking) and ChatGPT-5.5 (High) using a similar prompt (see Appendix), but uploading the three above-mentioned cleaned manually prepared CSV files. This time, ChatGPT reported rho values much lower than in the initial run, while Claude reported much higher values (Table 4).

Overall, this third task is the only task where a SOTA AI framework (specifically, Kosmos) demonstrated performance superior to that of a human expert. This level, however, is not yet achieved by general-purpose AI frameworks (Claude, ChatGPT).

## Discussion

This study compared two regimes of AI use on three scientific problems chosen to differ in domain and in the kind of evidence each requires. In the autonomous regime, an agentic AI framework received an open objective and executed without intervention; in the human-led regime, human experts fixed the scope, supplied curated data, and used AI as a coding and analysis assistant. Overall, autonomous operation failed on all three problems, except for one AI framework (Kosmos) on the third project, whereas the assisted regime produced reusable resources and modest but defensible performance (Table 1). Because the problems share little at the level of data or method, failures recurring across them point to general limitations of SOTA AI.

Several failure modes recurred independently of the domain. Data were silently omitted or substituted: Kosmos has not found published cohorts central to the resistance problem (TAURUS, UNITI-2), a serious problem where cohorts often hold only tens of patients, and on the third problem, ChatGPT-5.5 could not get molecular fingerprints and ended up building property-prediction models on experimental-protocol metadata. Required features could not be computed under execution-environment limits, confining Claude and ChatGPT to uncommon descriptors derivable from raw strings. Many candidate scores were presented without a single committed estimate, as in the Kosmos property report, inviting selection of the maximum and converting test scores into validation scores. Apparent success arose through unintended routes: in the audited recognition run, all correct matches for one paper came from PubChem name lookup rather than from reading the structures, with overall ground-truth agreement of only 17.8 percent.

Domain-specific failures were equally consequential. On the resistance problem, two near-identical cohorts from one research group (GSE16879 and GSE73661) were treated by Kosmos as independent test sets, inflating apparent generalization. For the recognition problem, MolVec, DECIMER, and MolScribe returned zero or near-zero precision and recall because they cannot abstain from images without molecules and cannot combine a scaffold with tabulated R-groups; the bottleneck is figure segmentation, not the recognition core. On property prediction, sensitivity to input noise yielded rho as high as 0.78 from non-structural columns.

Within human-defined scope, AI contributed substantively, accelerating the harmonization of nine studies into a modeling-ready table of 3256 samples with 967 labeled cases, feature computation, and code debugging. The assisted resistance models reached pre-treatment leave-one-cohort-out AUC near 0.63, post-treatment AUC of 0.75, and change-vector AUC averaging 0.84, and recovered expected and less-studied candidate genes. Property prediction was the single case in which an autonomous framework, Kosmos, exceeded the human expert, computing fingerprints, descriptors, and graph-neural-network embeddings. The Claude Code recognition run marks the boundary precisely: it owned environment setup and end-to-end execution with minimal supervision, yet ran the pipeline it was given and never reached for the workarounds an experienced chemist would apply, such as fragmenting a large molecule at its linker or adding macrocycle-aware detection. It automated the engineering, not the chemistry.

Our work has several significant limitations. The evaluation was qualitative and not designed to rank AI frameworks on numeric scores. The recognition benchmark on the second project was restricted to six heterogeneous papers (1 to 105 molecules each), so the reported precision ∼11% and recall ∼21% are approximate equal-weight averages. The property datasets contain only hundreds of molecules each. All results are tied to specific AI framework versions at one point in a rapidly changing field.

We conclude that current autonomous AI frameworks can carry out engineering and analysis once a competent human fixes the scope, supplies trustworthy data, and specifies the standard of evidence, but cannot yet be trusted to define a real problem, acquire and validate data, or engineer a novel solution without oversight. On real-life problems, as distinct from standard benchmarks, the productive use of AI in scientific research remains an art rather than a routine technology.

## Methods

### Evaluation philosophy

Evaluation was qualitative and practice-oriented. The aim was to identify approaches that work in practice rather than to rank AI systems on numeric scores. The standard of evidence was verification against computational artifacts and, where available, ground truth data. Intermediate narrative summaries of AI iterations were not treated as primary evidence, and every reported conclusion was checked against notebooks, generated files, statistical outputs, and visualizations produced within individual reasoning trajectories.

### Inflammatory treatment resistance

For preprocessing of the datasets, direct aggregation was infeasible because the cohorts differ across six axes: disease indication, therapeutic agent, biopsy site, profiling modality, temporal design, and endpoint definition. This CSV dataset was produced by a reproducible, version-controlled pipeline. A core script maps disparate inputs into a 21-column schema that records study identifier, anonymized patient identifier, sample identifier, timepoint, disease subtype, drug name and class, biopsy location, assay modality, binary response label, the precise response definition, the provenance of that definition, train and test split assignment, local data path, and curator notes.

A label-parsing module standardizes response annotations hidden in sample titles and metadata, and a feature module adds a biologically grounded 50-gene panel with explicit cross-platform mapping. Three choices preserve integrity: response definitions are retained as first-class fields, orthogonal boolean flags encode temporal relationships, and genes without unambiguous cross-platform mapping are encoded as missing rather than imputed. Initially, we attempted to preprocess nine relevant published datasets: CERTIFI, UNITI-2, RISK, GSE16879, GSE73661,TAURUS, Mennillo VDZ, R4RA, and STRAP. However, we found that CERTIFI and RISK have a large number of baseline timepoint rows, but lack response labels, which is why they could not be used by our prediction models. R4RA processed counts (not publicly available) were not obtained from QMUL due to temporal limitations of this project. Additionally, a within-patient change-vector matrix was constructed for the two longitudinal cohorts that contain paired baseline and post-treatment biopsies from the same patient, TAURUS (adalimumab, Crohn disease and ulcerative colitis) and the Mennillo vedolizumab subset (ulcerative colitis); these are the only cohorts in the precomputed table for which a post-minus-baseline vector is definable. The table was first filtered to these two cohorts, and multi-row visits, arising from several biopsy sites or cell groupings per visit in TAURUS and multiple imaging fields per visit in Mennillo, were collapsed by taking the mean of every numeric feature within each patient and timepoint, yielding one vector per visit. Patients retaining both a baseline and a post-treatment visit were kept, giving 41 paired patients (34 TAURUS and 7 Mennillo). The change vector was then computed as post minus baseline for every numeric feature, producing 80 delta features alongside 13 identifier and traceability columns.

Training, cross-validation, and testing ML models was performed in two nested loops, the outer loop going over various choices of the test set (either random or by cohort), and the inner loop for cross-validation for the remaining subsets (different in different iterations of the outer loop) to train ML models and optimize hyperparameters. This approach results in distributions of the test AUCs and numbers of relevant genes over different choices of the test set. For genes and pathways models, 100 different random test sets (each 25% of the whole dataset) were used to get distributions of the test AUC values and lists of important genes. The following ML models were tried: `LogReg_L2,RandomForest,GradBoosting, SVM_RBF, NeuralNet`. Per-study z-score normalization was performed. The best model and regularization coefficient (scanned in the range of 0.01 to 100) were chosen by 5-fold cross-validation. The delta dataset models were run over 100 different random test sets. Unless otherwise specified, L1 regularization was performed to simplify interpretation of the resulting models, because it provides a well-defined list of important features and, therefore, genes or pathways involved in inflammatory treatment resistance. Interpretability was assessed with SHAP values computed on held-out data.

#### Optical chemical structure recognition

Papers for our new benchmark were selected for recency, open-access availability, topic diversity, and image and rendering-style diversity. Figures were extracted in Docling ran in two variants: (1) with default settings, which resulted in the loss of multiple images with chemical structures, and (2) with settings tuned to recover small inline structures (using the heron layout backbone, with the picture-detection threshold lowered to 0.4 and page rasters rendered at images_scale 3.0) and a post-processing crop pass that retrieved individual table cells and filtered images below 5 kilobytes as noise. SMILES were canonicalized using `rdkit` (version 2026.3.2) with the default parameters of `rdkit.Chem.MolToSmiles`. Models were evaluated on exact matches of canonical SMILES (with stereo where defined); additionally, Tanimoto distances between molecules were computed on Morgan fingerprints and pairs with distances slightly less than 1 were manually checked for possible errors in SMILES processing or ground truth values. The evaluated ML model set spanned rule-based systems, deep-learning recognition models, and generalist vision-language models queried with a prompt requiring a SMILES-only output.

In the task of creating our benchmark, the limiting step was the construction of ground truth sets of SMILES drawn in the corresponding PDF files. Obtaining verified SMILES for every drawn structure proved difficult, in part because supplementary SMILES, when available, do not map to the structures actually drawn in figures. For example, Ref. ^86^ provided 10,806 SMILES in supplementary materials, but only 4 drawn structures in the main text, without specifying which SMILES were drawn. For this reason the current benchmark is temporarily restricted to six papers with SMILES sets fully curated by us (paper identifiers 1, 2, 4, 6, 16, 19), with the remaining papers to be completed in future work. Some extraction and recognition runs cover all 20 papers, but all quantitative results in the main text are reported on the above-listed six papers with curated ground truth SMILES sets. We purposefully refrain from disclosing the choice of the papers not included in the curated subset, because our intermediate results show that some of them may turn out to be not suitable for benchmarking for various nontrivial reasons, and we may replace them with other papers in the future.

To measure the performance, we use precision and recall defined in the common way, assuming the following interpretation:

- true positives are SMILES present both in the PDF file and in the AI output,
- false positives are SMILES absent from the PDF file, but present in the AI output,
- false negatives are SMILES present in the PDF file, but absent from the AI output.

(True negatives are not defined in this context. We count only exact coincidence of canonical SMILES; partial credits for “nearly correct” structures are not given.) Therefore, ***precision*** is the number of correct SMILES in the AI output divided by the total number of SMILES in the AI output, while ***recall*** is the number of correct SMILES in the AI output divided by the number of SMILES of molecules drawn in the PDF file.

#### Drug-design-related properties prediction

Three GDPsm1 datasets kindly provided by Ginkgo Datapoints were used: kinetic solubility (182 datapoints), microsomal stability (109 datapoints), and CYP2D6 inhibition (313 datapoints). Molecular structures were standardized with largest-fragment selection, neutralization, and canonical tautomer enumeration. Replicate assays were aggregated by median. Concentration-like targets were log-transformed, and censoring constraints were applied for the microsomal and inhibition endpoints. Featurization combined Morgan fingerprints, a physicochemical descriptor block filtered for nonzero variance, and a ChemBERTa embedding compressed by principal component analysis. In total, about 1800 descriptors were computed. Scaffold-based splitting using Bemis-Murcko scaffolds was used to select the test subset, simulating prospective lead optimization scenario. ML models spanned Ridge Regression, ElasticNet, Random Forest, and Gradient Boosting. Spearman rank correlation was the primary metric, matching deployment criteria in which ranking compounds, e.g. for synthesis prioritization, takes precedence over absolute numeric error of predictions.

## Declaration on AI use in this work

AI frameworks, including large language models, were used as coding and debugging assistants and as first-draft analysts throughout the design, implementation, and analysis stages. In the engineering loop, AI agents were used as described above to try to run automated scientific research. In the document loop, AI assisted in organizing dataset metadata and structuring technical reports. All quantitative claims reported here were verified by human authors against the underlying computational artifacts.

## Acknowledgements

We are deeply grateful to Dr. Jacqueline Valeri for valuable discussions on the computer vision for data mining project. We would also like to thank Ginkgo Datapoints for providing access to the GDPsm1 datasets, and the authors of Kosmos for free credits to students and academics to run their AI framework. J.L. was supported by an NIH R01 award (GM143370) and I.E.L. by the College of Pharmacy Summer Undergraduate Research Fellowship (SURF) at Purdue University.

## Appendix

### Inflammatory treatment resistance

Initial prompt: “Using relevant, publicly available datasets, identify predictors of treatment non-response in immune-mediated inflammatory diseases. Interpret these results in terms of molecular mechanisms of diseases and genes involved.”

Kosmos: https://platform.edisonscientific.com/kosmos/602c81bb-e622-4c9f-a1ff-d5b043d093ff

Claude run A: https://claude.ai/share/e481e469-bb3f-426a-99e1-ceef5cedbf70

Claude run B: https://claude.ai/share/506a8b3a-ce44-438e-ab0c-06b5bd84bc95

ChatGPT: https://chatgpt.com/share/6a398ee3-0438-83ea-8b6c-a55ec01eb930

Second prompt: “Using the uploaded dataset, identify predictors of treatment non-response in immune-mediated inflammatory diseases. Interpret these predictors in terms of molecular mechanisms of disease and the genes involved.” (preprocessed CSV file attached)

Kosmos: https://platform.edisonscientific.com/kosmos/6c688829-b68a-418e-bddc-eeb3c41e3be0

Claude: https://claude.ai/share/3cb830d1-2714-4361-843e-dc0282cbbf00

### Computer vision on organic molecules

Prompt: “Write Python code that takes a PDF file of a scientific paper as input and recognizes all molecules drawn in images within that PDF file. The code should produce a CSV file with two columns: the molecule label used in the paper, which may be an empirical name or a shorthand notation such as “2” or “9c”, and the corresponding SMILES. The SMILES must be recognized directly from the images, not looked up from the names and not extracted from the text of the PDF. The code should handle real papers that have complex layouts, including figures, schemes, and tables that may each contain multiple molecules, a single molecule, or no molecules. It should also handle cases where a core scaffold is shown together with several R-group variants rather than a full structure for each molecule, and molecules drawn in various styles.”

Claude: https://claude.ai/share/82f33940-ba56-4a31-b7a8-796cda6e7444 ChatGPT: https://chatgpt.com/share/6a384706-2fc0-83ea-a7af-e2c82e17138a

### Prediction of drug-design-related properties

Initial prompt: “Using the uploaded datasets, build machine learning models for predicting kinetic solubility, microsomal stability, and P450 inhibition. Measure the performance by test-set Spearman rho values.” (three original XLSX files attached)

Second prompt: “Using the uploaded cleaned Ginkgo ADMET CSV files, automatically build and evaluate ML models for each ADMET target using the available descriptors and embeddings. Report the preprocessing steps, train/test split, selected models, and Spearman rho results. Do not assume missing information; inspect the files first and make the results checkable.” (three pre-processed CSV files attached)

Links to Claude, ChatGPT, and Kosmos runs are available by request to prevent uncontrollable leakage of the datasets provided by Ginkgo Datapoints.

